# cTAGE5 acts as a Sar1 GTPase regulator for collagen export

**DOI:** 10.1101/452904

**Authors:** Norito Sasaki, Masano Shiraiwa, Miharu Maeda, Tomohiro Yorimitsu, Ken Sato, Toshiaki Katada, Kota Saito

## Abstract

**Secretory proteins synthesized within the endoplasmic reticulum (ER) are exported via coat protein complex II (COPII)-coated vesicles. The formation of the COPII-coated vesicles is initiated by activation of the small GTPase, Sar1. cTAGE5 directly interacts with a guanine-nucleotide exchange factor (GEF), Sec12, and a GTPase-activating protein (GAP) of Sar1, Sec23. We have previously shown that cTAGE5 recruits Sec12 to the ER exit sites for efficient production of activated Sar1 for collagen secretion. However, the functional significance of the interaction between cTAGE5 and Sec23 has not been fully elucidated. In this study, we showed that cTAGE5 enhances the GAP activity of Sec23 toward Sar1. In addition, the interaction of cTAGE5 with Sec23 is necessary for collagen exit from the ER. Our data suggests that cTAGE5 acts as a Sar1 GTPase regulator for collagen secretion.**

## INTRODUCTION

Secretory proteins synthesized within the endoplasmic reticulum (ER) are exported via coat protein complex II (COPII)-coated vesicles (Miller and Schekman, 2013). COPII vesicle formation is initiated by the activation of small GTPase, Sar1, by its guanine-nucleotide exchange factor (GEF), Sec12 (Nakano and Muramatsu, 1989; Barlowe and Schekman, 1993). The N-terminus amphipathic helix of activated Sar1 penetrates the ER membranes and induces membrane curvature (Lee *et al.*, 2005; Long *et al.*, 2010; Settles *et al.*, 2010). Sar1 then forms the pre-budding complex with Sec23/Sec24, the inner coat complex. Sec24 interacts with cargo receptors to recruit cargoes into nascent vesicles (Matsuoka *et al.*, 1998; Bi *et al.*, 2002; Sato and Nakano, 2005). Finally, Sec13/Sec31, the outer coat complex, enhances the GAP activity of Sec23, inducing Sar1 GTP hydrolysis, and completes vesicle formation (Yoshihisa *et al.*, 1993; Antonny *et al.*, 2001; Bi *et al.*, 2007). Sec16, another peripheral membrane protein, is also involved in COPII vesicle formation by modulating Sar1 GTPase activity (Kung *et al.*, 2012; Yorimitsu and Sato, 2012).

Mammalian cells export various cargoes, including bulky molecules, such as collagens (Saito and Katada, 2015). It has been debated that how collagens, which form larger than 300 nm-long rigid structures are exported from the ER, since conventional COPII-coated vesicles are 60–90 nm in diameter (Fromme and Schekman, 2005; Malhotra and Erlmann, 2015). Recently, collagens were reported to be packaged into COPII-coated large spherical structures (Gorur *et al.*, 2017), and the mechanisms underlying extra-sized COPII vesicle formation are emerging (Venditti *et al.*, 2012; Nogueira *et al.*, 2014; Santos *et al.*, 2015; Raote *et al.*, 2017). We have identified TANGO1 as a cargo receptor for collagens, and revealed that TANGO1 forms membrane spanning macromolecular complex with multiple cTAGE5 and Sec12 molecules at ER exit sites (Saito *et al.*, 2009; Maeda *et al.*, 2016).

We previously reported that cTAGE5 recruits Sec12 to the ER exit sites (Saito *et al.*, 2014). Our data suggested that cTAGE5 does not affect GEF activity of Sec12 toward Sar1 (Saito *et al.*, 2014), but the concentration of Sec12 at ER exit sites accounts for the efficient production of activated Sar1 required for collagen secretion (Tanabe *et al.*, 2016; Saito *et al.*, 2017). On the contrary, we also reported that proline-rich domain (PRD) of both cTAGE5 and TANGO1 interact with Sec23 and Sec24 by yeast two-hybrid analysis (Saito *et al.*, 2009; Saito *et al.*, 2011), but the functional significance of these interactions has not been investigated.

In the present study, we revealed that cTAGE5 enhances the GAP activity of Sec23 towards Sar1. In addition, the interaction of cTAGE5 and Sec23 is required for collagen exit from the ER. We proposed a model that cTAGE5 acts as a Sar1 GTPase regulator for collagen secretion.

## RESULTS AND DISCUSSION

### cTAGE5 enhances the GAP activity of Sec23/Sec24 toward Sar1

We have previously shown that cTAGE5 interacts with Sec23/Sec24 (Saito *et al.*, 2011); however, whether the interaction with cTAGE5 has any influence on the GAP activity of Sec23 toward Sar1 has not yet been investigated. We measured the GTPase activity of Sar1 by Pi release on the liposome in the presence of Sec23/Sec24 and cTAGE5 (Figure S1). We purified the cytoplasmic domain of cTAGE5 (61–804 aa) with His_6_ tag on the N-terminus, so that it can be recruited to DOGS-NTA-containing liposome in the topology same as that of the physiological ER membranes (Cabrera *et al.*, 2014; Ebine *et al.*, 2014). As shown in Figure 1A in columns 5 and 6, released Pi due to the action of Sar1 GTPase was increased with the addition of Sec23/Sec24, confirming that Sec23/Sec24 complex acts as a GAP for Sar1. Interestingly, when His_6_-tagged cTAGE5 cytoplasmic domain was introduced into the DOGS-NTA-containing liposome, released Pi was significantly upregulated in the presence of Sec23/Sec24 (Figure 1A, columns 6 and 9). However, cTAGE5 without Sec23/Sec24 has no stimulating activity toward Sar1 GTPase (Figure 1A, columns 4, 5, and 8), indicating that cTAGE5 does not act as a GAP by itself, but enhances the GAP activity of Sec23 toward Sar1. Although the addition of His_6_-tagged Sec12 cytoplasmic region (1–386 aa) alone has little effects on the production of free Pi by Sar1 (Figure 1A, column 5 and 7), the combination with Sec23/Sec24 significantly increased the amount of released Pi (Figure 1A, column 11), consistent with the previous finding in yeast (Barlowe and Schekman, 1993). Moreover, further addition of cTAGE5 increases the production of free Pi (Figure 1A, columns 11 and 12), confirming that cTAGE5 enhances the GAP activity of Sec23 toward Sar1. Next, to check the specificity of cTAGE5, we performed the assay with liposome free of DOGS-NTA, so that the cTAGE5 is not attached to the liposome in this condition. As shown in Figure 1B, the addition of cTAGE5 failed to enhance the GAP activity of Sec23 in this condition. In contrast, cTAGE5 efficiently enhances the GAP activity of Sec23 toward Sar1 on the liposome with DOGS-NTA in a concentration-dependent manner (Figure 1C). As shown in Figure S2, the activity of cTAGE5 is statistically significant. Interestingly, cTAGE5 enhances the GAP activity of Sec23 more efficiently than Sec13/31 does (Please refer to the concentration of Figure 1C and D). It is probably because cTAGE5 is attached to the membrane so that local concentration of cTAGE5 is higher than that of Sec13/31 around membranes.

**Figure 1.**
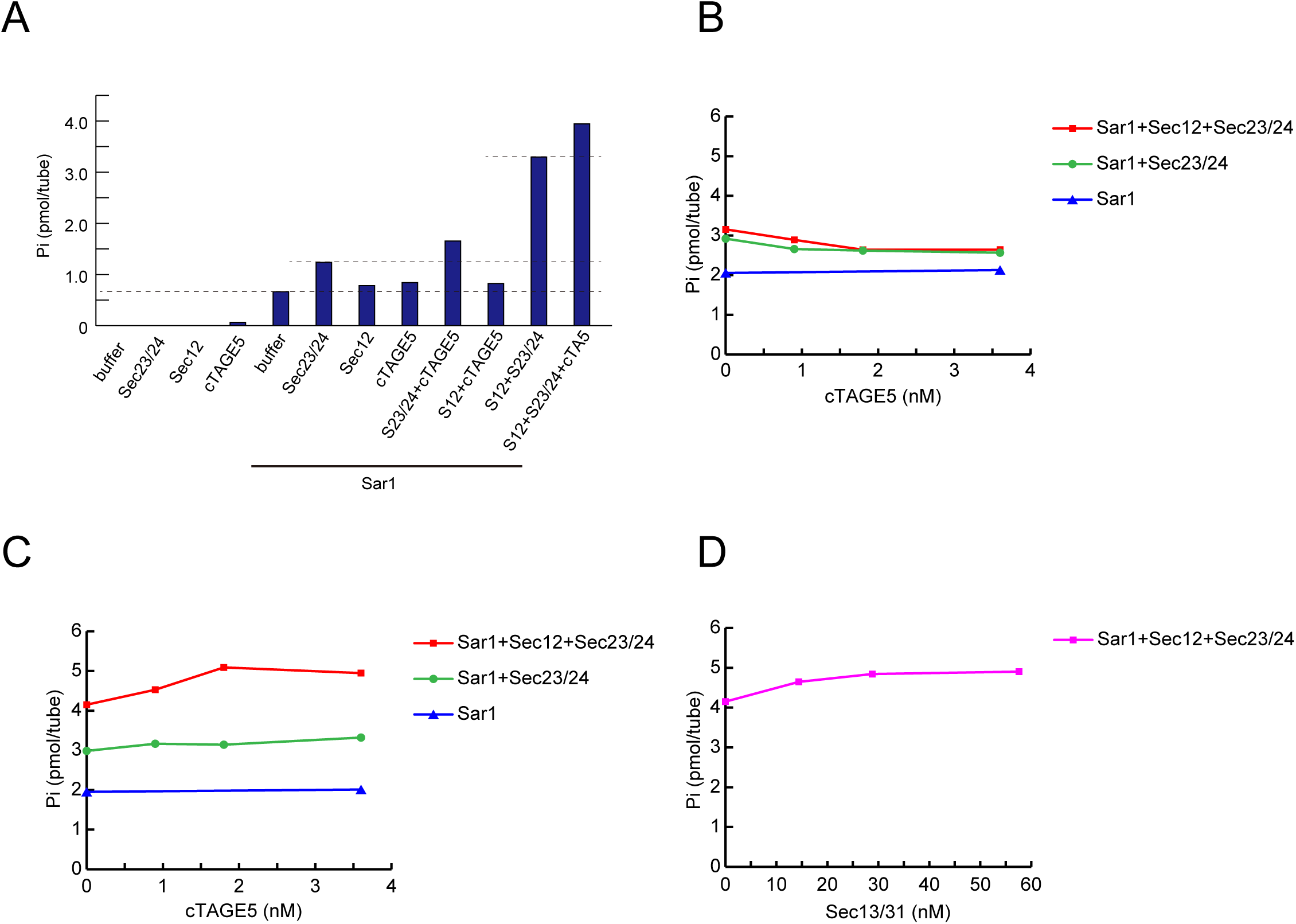
cTAGE5 enhances the GAP activity of Sec23 toward Sar1. (A) 200 nM Sar1 was incubated with a buffer consisting of 25 *µ*g/ml Ni liposome (20%), 20 mM HEPES-KOH (pH 7.4), 100 mM KOAc, 6 mM Tris-HCl, 45 mM NaCl, 1 mM MgCl_2_, 1.6% Glycerol, 133 nM GDP, 2 *µ*M [γ-^32^P] GTP, 50 *µ*M AppNHp, 67 *µ*M β-ME, 67 *µ*M AEBSF, in the presence or absence of 8 nM FLAG-Sec12 (1–386 aa)-His_6_, 15 nM HA-Sec23A/FLAG-Sec24D, 3.6 nM cTAGE5 for 1 h at 30°C. The amount of free ^32^Pi was quantified. (B) 200 nM Sar1 was incubated with a buffer consisting of 25 *µ*g/ml liposome without Ni, 20 mM HEPES-KOH (pH 7.4), 100 mM KOAc, 6 mM Tris-HCl, 45 mM NaCl, 1 mM MgCl_2_, 1.6% Glycerol, 133 nM GDP, 2 *µ*M [γ-^32^P] GTP, 50 *µ*M AppNHp, 67 *µ*M β-ME, 67 *µ*M AEBSF, in the presence or absence of 8 nM FLAG-Sec12 (1–386 aa)-His_6_, 15 nM HA-Sec23A/FLAG-Sec24D and the indicated concentration of cTAGE5 for 1 h at 30°C. The amount of free ^32^Pi was quantified. (C, D) 200 nM Sar1 was incubated with a buffer consisting of 25 *µ*g/ml Ni liposome (20%), 20 mM HEPES-KOH (pH 7.4), 100 mM KOAc, 6 mM Tris-HCl, 45 mM NaCl, 1 mM MgCl_2_, 1.6% Glycerol, 133 nM GDP, 2 *µ*M [γ-^32^P] GTP, 50 *µ*M AppNHp, 67 *µ*M β-ME, 67 *µ*M AEBSF, in the presence or absence of 8 nM FLAG-Sec12 (1–386 aa)-His_6_, 15 nM HA-Sec23A/FLAG-Sec24D and the indicated concentration of (C) cTAGE5 or (D) Sec13/31 for 1 h at 30°C. The amount of free ^32^Pi was quantified.

### Construction of cTAGE5 point mutants lacking Sec23A-binding activity

We have previously shown that PRD of both cTAGE5 and TANGO1 interact with Sec23/Sec24 by yeast two-hybrid assay (Saito *et al.*, 2009; Saito *et al.*, 2011). To further analyze these interactions, we utilized recombinant proteins and checked whether they directly interact by *in vitro* binding assay. Although PRD of both cTAGE5 and TANGO1 efficiently interact with Sec23, they failed to bind Sec24 (Figure 2A). We concluded that cTAGE5 and TANGO1 directly bind to Sec23, but not with Sec24. We then screened cTAGE5 mutant incapable of binding to Sec23. We employed mutations in cTAGE5 PRD by error-prone PCRs and isolated clones, which exhibited reduced interaction with Sec23 by yeast two-hybrid assay. Most of the positive clones contained non-sense mutations; however, we identified one missense mutation, R757G (Figure 2B and C).

**Figure 2.**
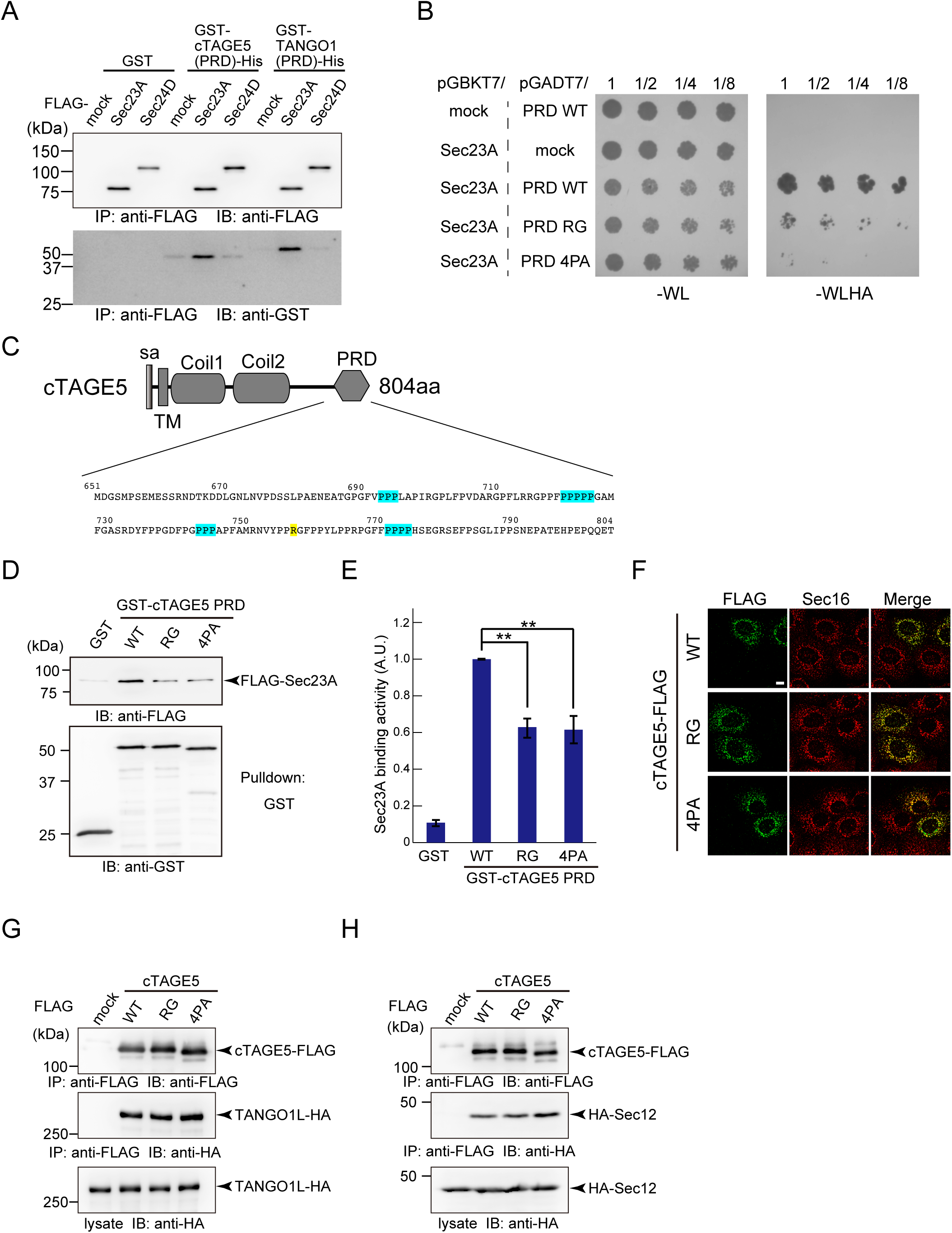
Construction of cTAGE5 mutants lacking Sec23-binding activity. (A) Purified FLAG-Sec23A or FLAG-Sec24D was immobilized onto FLAG agarose beads and incubated with recombinant GST or GST-tagged cTAGE5-PRD (651–804aa)-His_6_ or GST-tagged TANGO1-PRD (1651–1907 aa)-His_6_. Beads were washed and eluted with FLAG peptide. Eluted proteins were subjected to SDS-PAGE followed by western blotting with anti-FLAG and anti-GST antibodies. (B) PRD regions of cTAGE5 mutants in pGADT7 plasmids were co-transformed with pGBKT7 plasmids containing Sec23A into AH109 yeast strains and grown on tryptophan-, leucine-deficient plate (-WL). Interactions were investigated by observing the cell growth on tryptophan-, leucine-, histidine-, and adenine-deficient plate (-WLHA). (C) Schematic representation of human cTAGE5 domain organization. sa, signal anchor; TM, transmembrane; Coil, coiled-coil domain; PRD, proline-rich domain. The position of R757 is shaded in yellow and the positions of 4PA mutations are shaded in light blue. (D) Recombinant GST or GST-tagged cTAGE5-PRD (651–804aa)-His_6_ wild-type or cTAGE5-PRD R757G or cTAGE5-PRD 4PA were immobilized to glutathione sepharose resin and incubated with FLAG-Sec23A. Resins were washed and eluted with glutathione. Eluted proteins were subjected to SDS-PAGE followed by western blotting with anti-FLAG and anti-GST antibodies. (E) Quantification of Sec23A immunoblots (*n* = 9). The band intensities were normalized to those of GST blots. Error bars represent mean ± SEM. \*\**P* < 0.001. (F) HSC-1 cells were treated with cTAGE5 siRNA and cultured for 24 h. Then, FLAG-tagged cTAGE5 wild-type, cTAGE5 R757G, or cTAGE5 4PA were transfected and further cultured for 24 h. The cells were fixed and stained with Sec16 and FLAG antibodies. Bars = 10 *µ*m. (G, H) 293T cells were transfected with FLAG tag only, FLAG-tagged cTAGE5 wild-type, cTAGE5 R757G, or cTAGE5 4PA with HA-tagged (G) TANGO1L or (H) Sec12. Cell lysates were immunoprecipitated with anti-FLAG antibody and eluted with FLAG peptide. Eluates and cell lysates were analyzed by SDS-PAGE followed by western blotting with FLAG or HA antibodies.

During our analysis, Ma et al. reported that PPP motifs in the PRD regions of both cTAGE5 and TANGO1 are responsible for the interaction with Sec23A (Ma and Goldberg, 2016). Thus, we also prepared 4PA mutant, in which four regions of consecutive prolines were changed to alanines (a total of 15 prolines were changed to alanines) (Ma and Goldberg, 2016) (Figure 2C). As shown in Figure 2B, the 4PA mutant was also incapable of interacting with Sec23 in the yeast two-hybrid assay. Next, we checked the interaction by *in vitro* binding assay. Both RG and 4PA mutants showed reduced binding to Sec23, consistent with the yeast two-hybrid assay (Figure 2D and E).

Then, we examined whether mutants correctly localized to the ER exit sites. As shown in Figure 2F, both RG and 4PA mutants extensively co-localized with Sec16, a bone-fide ER exit site marker, suggesting that the mutations do not affect its localization. We previously reported that the localization of the ER exit sites within the ER is defined by the interaction between TANGO1 and Sec16, and cTAGE5 is recruited to the ER exit sites by the interaction with TANGO1 (Maeda *et al.*, 2017). Current data further supported the idea that cTAGE5 is not likely to be localized to the ER exit sites by the interaction with Sec23/Sec24.

Next, we checked whether cTAGE5 mutants retain the properties to interact with TANGO1 and Sec12. As shown in Figure 2G and H, both the mutants were still bound to TANGO1L and Sec12, indicating that the mutations do not destroy the overall conformation of cTAGE5, except for its affinity to Sec23.

### PRD of cTAGE5 is responsible for enhancing the GAP activity of Sec23

Because PRD of cTAGE5 is responsible for interacting with Sec23, we speculated that this domain is also involved in the GAP enhancing activity against Sec23. Thus, we made recombinant PRD of cTAGE5 and checked the activity against Sec23. As shown in Figures 3 and S3, the addition of cTAGE5 PRD into DOGS-NTA liposome with Sar1, Sec12, and Sec23/Sec24 significantly enhanced the Pi release in a concentration-dependent manner. Conversely, the addition of GST has no effects on the free Pi production (Figures 3 and S3), indicating that cTAGE5 PRD has a property to enhance the GAP activity of Sec23 toward Sar1. Next, we examined the effects of cTAGE5 mutants with reduced Sec23-binding on the GAP enhancing activity. Interestingly, although RG mutant retained the activity at the normal level, 4PA mutant failed to enhance the GAP activity of Sec23 toward Sar1 (Figures 3 and S3). These results indicated that the domain responsible for GAP-enhancing activity and Sec23-binding could be separable within the PRD of cTAGE5.

**Figure 3.**
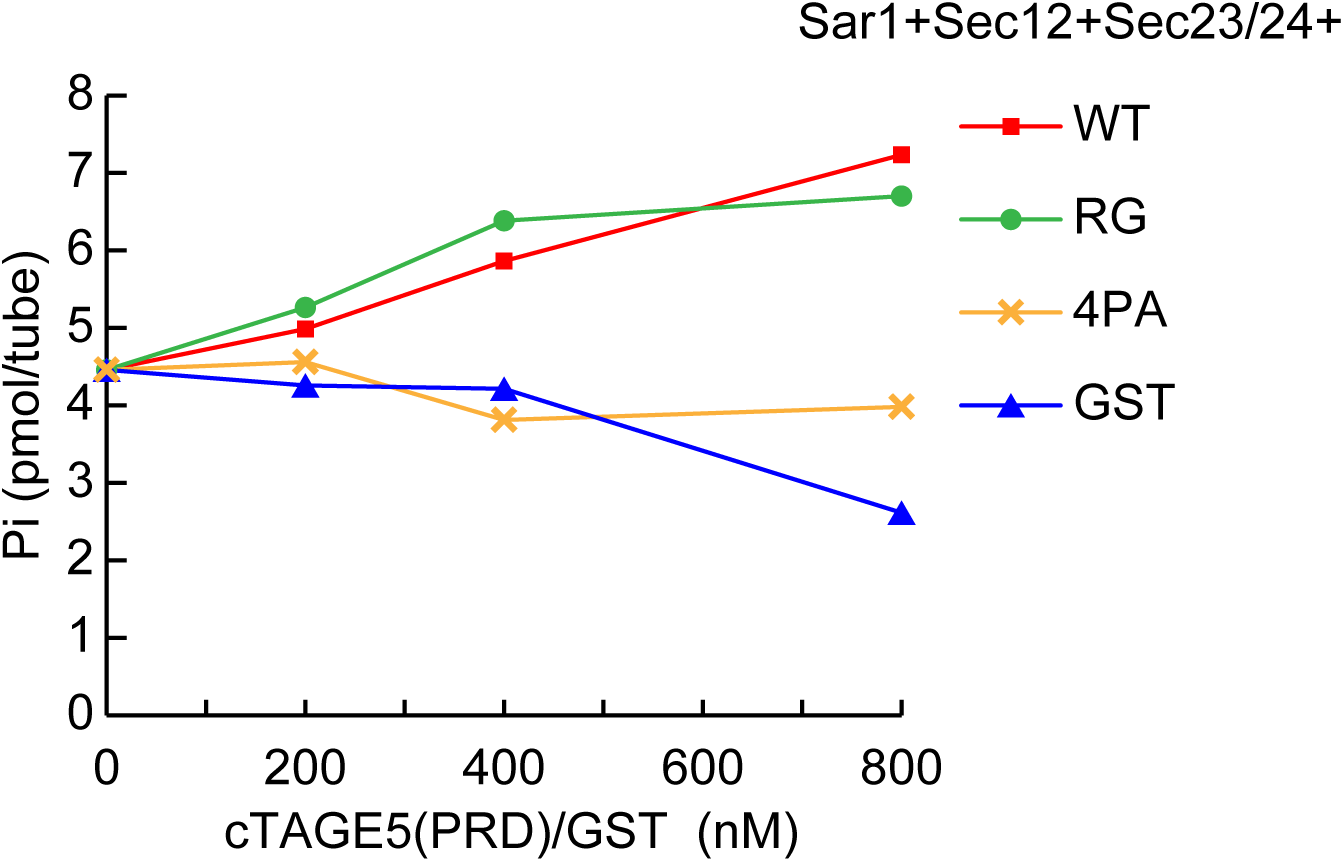
PRD of cTAGE5 enhances GAP acitivity of Sec23 toward Sar1. 200 nM Sar1 was incubated with a buffer consisting of 25 *µ*g/ml Ni liposome (10%), 20 mM HEPES-KOH (pH 7.4), 100 mM KOAc, 8 mM Tris-HCl, 60 mM NaCl, 1 mM MgCl_2_, 1.6% Glycerol, 133 nM GDP, 2 *µ*M [γ-^32^P] GTP, 50 *µ*M AppNHp, 67 *µ*M β-ME, 67 *µ*M AEBSF, 8 nM FLAG-Sec12 (1–386 aa)-His_6_, 15 nM HA-Sec23A/FLAG-Sec24D and the indicated concentration of His_6_-GST, His_6_-cTAGE5 (651–804 aa)-GST wild-type, R757G, 4PA for 1 h at 30°C. The amount of free ^32^Pi was quantified.

### cTAGE5 with reduced Sec23-binding activity failed to secrete collagen VII from the ER

Next, we examined whether the mutants could promote collagen VII secretion from the ER. As previously reported (Saito *et al.*, 2014; Maeda *et al.*, 2016; Tanabe *et al.*, 2016), we quantified the signals of accumulated collagen VII within the ER as an index of its secretion. The expression of both the mutants with reduced Sec23-binding activity to the ER exit sites could not rescue the block of collagen VII secretion induced by cTAGE5 knockdown, although the wild-type cTAGE5 rescued collagen secretion (Figure 4).

**Figure 4.**
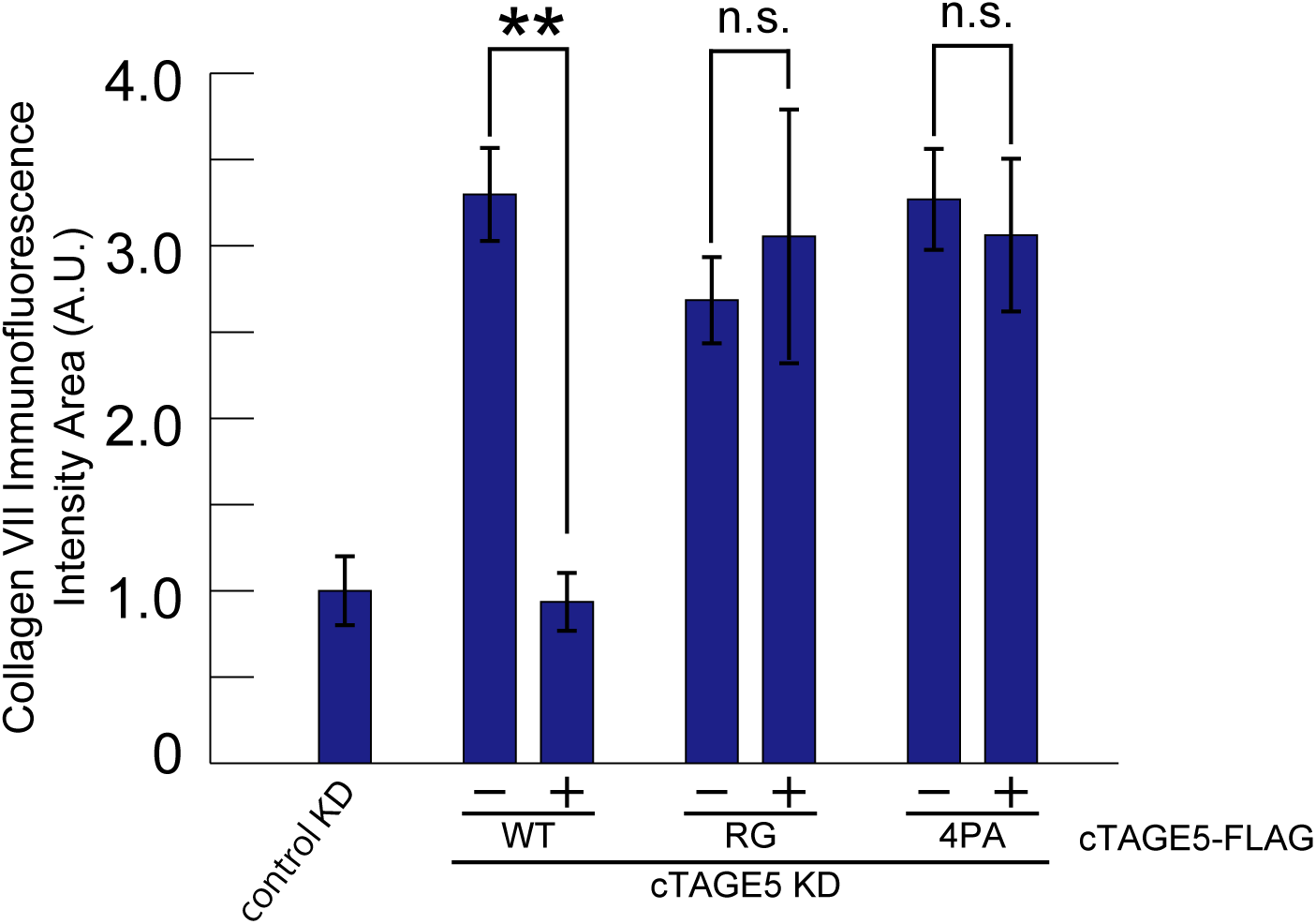
cTAGE5 mutants with reduced Sec23-binding activity failed to secrete collagen VII. HSC-1 cells were treated with control or cTAGE5 siRNA and cultured for 24 h. For cTAGE5 siRNA-treated cells, FLAG-tagged cTAGE5 wild-type, cTAGE5 R757G or cTAGE5 4PA were transfected and further cultured for 24 h. The cells were fixed and stained with collagen VII and FLAG antibodies. Collagen VII immunofluorescence signal per cell (arbitrary units, A.U.) were quantified in each cell category described below. The cells positively stained with FLAG antibody were categorized as the constructs expressed, and the surrounding unstained cells were categorized as non-transfected counterparts. Within each well, cells transfected with constructs are labeled as + and non-transfected cells are labeled as −. Analysis of variance. Error bars represent mean ± SEM; \*\**P* < 0.001; n.s., *P* > 0.05. The data shown are from single representative experiment out of four replicates. Cells treated with control siRNA (*n* = 65); cells treated with cTAGE5 siRNA and wild-type- (*n* = 162); wild-type + (*n* = 53); R757G − (*n* = 151); R757G + (*n* = 53); 4PA − (*n* = 177); 4PA + (*n* = 57).

### cTAGE5 acts as a Sar1 GTPase regulator for collagen export

cTAGE5 is an integral membrane protein containing two coiled coil domains and PRD in the cytoplasmic region. cTAGE5 belongs to the cTAGE gene family, consisting of 6 genes coding for highly homologous proteins and 9 pseudogenes according to the HUGO Gene Nomenclature Committee (Comtesse *et al.*, 2001). Until now, we have been focusing on studying cTAGE5, and the possibility that the antibodies we used might cross-react with other cTAGE-family proteins cannot be ruled out. However, this seems unlikely because the siRNAs that we used efficiently depleted proteins recognized by cTAGE5 antibodies, and we revealed these siRNAs are specific for cTAGE5, and not cross-reactive with the other five proteins, by analyzing RefSeq database (Saito *et al.*, 2011; Saito *et al.*, 2014; Tanabe *et al.*, 2016). Thus, we believe that our study describes the results only for cTAGE5, and the influence on other cTAGE family members would be very limited. Recently, cTAGE5 has been reported to be involved in collagen, VLDL, and insulin secretion (Saito *et al.*, 2011; Saito *et al.*, 2014; Santos *et al.*, 2016; Tanabe *et al.*, 2016; Wang *et al.*, 2016; Fan *et al.*, 2017). We previously reported that cTAGE5 directly interacts with Sec12 via cytoplasmic region just after the membrane-spanning domain and this interaction is specifically required for collagen secretion (Saito *et al.*, 2014; Tanabe *et al.*, 2016). In addition, we revealed that the second coiled coil domain of cTAGE5 is responsible for interacting with TANGO1, a collagen cargo receptor (Saito *et al.*, 2011), and cTAGE5 can form homo-multimer (Maeda *et al.*, 2016).

In this study, we revealed that the PRD of cTAGE5 is not only responsible for Sec23 binding, but also for the activation of Sec23 GAP activity toward Sar1. Thus, we proposed that TANGO1 recruits cTAGE5 multimer for regulation of Sar1 GTPase in the vicinity of the ER exit sites. Sec12, concentrated around ER exit sites via interaction with cTAGE5, efficiently produces the activated Sar1 around ER exit sites. Sar1, then, might be involved in the collagen-containing tubule formation (See the discussion blow), and is efficiently hydrolyzed by Sec23, the activity of which is enhanced by the interaction with cTAGE5 or Sec31. This hydrolysis of GTP by Sar1 might be important for completing the collagen-containing carrier formation.

Notably, in the *in vitro* assay, liposomes incubated with the GTP-restricted form of Sar1 mutant (Sar1 H79G) or with Sar1 and non-hydrolyzable GTP analogs, such as GMP-PNP and GTPγS, induced tubules long enough to accommodate collagens inside (Long *et al.*, 2010; Bacia *et al.*, 2011). These tubular structures still attach to the liposome, implying that GTP hydrolysis by Sar1 is necessary for carriers to detach from liposomes. Moreover, cryo-electron microscopy analysis revealed that giant unilamellar vesicles incubated with non-hydrolyzable Sar1, Sec23/Sec24, and Sec13/Sec31 produce tubes coated with Sec23/Sec24 and Sec13/Sec31. Interestingly, the predicted model indicates that tubule coated with Sec23/Sec24 recruits less Sec13/Sec31 than the spherical vesicles do (Zanetti *et al.*, 2013). Thus, it is interesting to speculate that cTAGE5-mediated activation of Sec23 GAP activity might be important only for completing the formation of large cargo carriers from the tubular structures.

On the contrary, Sec31 has also been reported to be involved in collagen secretion. Ubiquitylation of Sec31 regulated by calcium-binding proteins and Cul3-KLHL12 leads to the large carrier formation (Jin *et al.*, 2012; McGourty *et al.*, 2016). It has not been investigated whether ubiquitylation of Sec31 has any effects on the activity toward Sec23. The mechanism underlying the coordination between the GAP-enhancing activities of Sec31 and cTAGE5 awaits further investigation.

Interestingly, RG mutant, which retains the property to enhance the Sec23 GAP activity, but exhibits reduced binding to Sec23, fails to secrete collagen VII from the ER. These data imply that the regions of cTAGE5 responsible for GAP enhancing activity and interaction with Sec23 could be separable within the cTAGE5 PRD. Alternatively, if the same regions are responsible for the activity and interaction, it suggests that RG mutant has a higher GAP enhancing activity than that of wild-type cTAGE5. It also suggested that interaction of cTAGE5 with Sec23 is necessary for collagen secretion, in addition to enhancing the GAP activity of Sec23. How this interaction participates in collagen secretion needs to be revealed in the future studies. In this study, we revealed that cTAGE5 enhances the GAP activity of Sec23 toward Sar1. In addition, the interaction of cTAGE5 and Sec23 is necessary for collagen exit from the ER. Thus, cTAGE5 acts as a Sar1 GTPase regulator for collagen secretion.

## MATERIALS AND METHODS

### Antibodies

Anti-collagen VII monoclonal antibody (NP-185) was kindly provided by Dr. Lynn Sakai. Other antibodies were used as described previously (Maeda *et al.*, 2017).

### Constructs

cTAGE5 rescue constructs were made as described previously (Saito *et al.*, 2014; Maeda *et al.*, 2017). cTAGE5-4PA construct was made by introducing following mutations (P693A P694A P695A P720A P721A P722A P723A P724A P743A P744A P745A P771A P772A P773A P774A).

### Cell culture and transfection

HeLa, HSC-1, and 293T cells were cultured in DMEM supplemented with 10% fetal bovine serum. Lipofectamine RNAi max (Thermofisher) was used for transfecting siRNA. For plasmids transfection, polyethylenimine “MAX” (polysciences) or FuGENE 6 (Promega) were used.

### Recombinant human protein purification

Proteins used for GTPase hydrolysis assay were all from humans. Baculovirus encoding FLAG-Sec12 (1–386 aa)-His_6_, His_6_-cTAGE5 (61–804 aa)-FLAG was made with Bac-to-Bac Baculovirus Expression System according to manufacturer’s protocol (Life Technologies). Sf9 cells infected with virus were collected. Each protein was purified with FLAG M2 agarose beads (Sigma-Aldrich). Elution was made with FLAG peptide, then the buffers were exchanged with 20 mM HEPES-KOH (pH 7.4), 160 mM KOAc, 1 mM MgCl_2_ by desalting column. Sec13/FLAG-Sec31a was cloned into pFastBacDual vector and baculovirus was produced. Protein was purified from infected Sf9 cells with FLAG M2 agarose beads (Sigma-Aldrich) followed by elution with FLAG peptide. The buffer was exchanged with TBS/5% (w/v) glycerol by desalting column. FLAG-Sec23A, FLAG-Sec24D were expressed in 293T cells and purified with FLAG M2 agarose beads (Sigma-Aldrich). Elution was made with FLAG peptide. The buffer was exchanged with TBS/0.05% Lubrol-PX by desalting column. HA-Sec23/FLAG-Sec24 were expressed in 293T cells and purified with FLAG M2 agarose beads (Sigma-Aldrich). Elution was made with FLAG peptide. The buffer was exchanged with TBS/5% (w/v) glycerol by desalting column. GST-Sar1a was expressed in *Escherichia coli* and purified with glutathione sepharose (GE Healthcare). Then GST tag was cleaved by thrombin protease followed by dialysis with 20 mM HEPES-KOH (pH 6.8), 160 mM KOAc, 5 mM MgCl_2_, 5 mM β-ME, 0.5 mM AEBSF, 10 *µ*M GDP, 5% (w/v) glycerol. His_6_-cTAGE5 (651–804 aa)-GST wild-type, His_6_-cTAGE5 (651–804 aa)-GST R757G, His_6_-cTAGE5 (651–804 aa)-GST 4PA were expressed in *Escherichia coli* and purified with glutathione sepharose (GE Healthcare). The GST tags were cleaved by prescission protease and further purified with Ni sepharose 6 Fast Flow (GE Healthcare) and eluted with imidazole. The buffers were exchanged with TBS by desalting columns. His_6_-GST expressed in *Escherichia* was purified by Ni sepharose 6 Fast Flow and eluted with imidazole. The buffer was exchanged with TBS by desalting column.

### Liposome Preparation

Lipids were purchased from Avanti Polar Lipids, except sphingomyelin (Enzo life science) and cholesterol (nakalai tesque). The lipid mixture was evaporated and then resuspended in 20 mM Hepes-KOH, pH 7.4, 160 mM KOAc, 1 mM MgCl_2_ followed by sonication. Liposome without Ni consists of di-oleoyl-phosphatidylcholine (DOPC; 54%), di-oleoyl-phosphatidyl-ethanolamine (DOPE; 21%), soy phosphatidylinositol (PI; 9%), cholesterol (Cho; 7%), di-oleoyl-phosphatidyl-serine (DOPS; 3%), sphingomyelin (SM; 3%), cystidine diphosphate diacylglycerol (CDP-DAG; 2%), dioleoyl-phosphatidic acid (DOPA; 1%). Liposome with Ni (20%) consists of DOPC; 41%, DOPE; 18%, PI; 5%, Cho; 7%, DOPS; 3%, SM; 3%, CDP-DAG; 2%, DOPA; 1%, and 1,2-Dioleoyl-sn-Glycero-3-[(N-(5-amino-1-carboxypentyl)iminodiacetic acid)succinyl](nickel salt)(DGS-NTA; 20%). Liposome with Ni (10%) consists of DOPC; 48.6%, DOPE; 18.9%, PI; 8.1%, Cho; 6.3%, DOPS; 2.7%, SM; 2.7%, CDP-DAG; 1.8%, DOPA; 0.9%, and DGS-NTA; 10%.

### GTP hydrolysis assay

His_6_-cTAGE5 (61–804 aa)-FLAG, FLAG-Sec12 (1–386 aa)-His_6_ and liposome were preincubated in a buffer consisting of 20 mM Hepes-KOH, pH 7.4, 160 mM KOAc, 1 mM MgCl_2_ for 2h at 4°C. Then, Sar1 GTP hydrolysis were initiated in a buffer consisting of 25 *µ*g/ml of liposome, 20 mM HEPES-KOH (pH 7.4), 100 mM KOAc, 6 mM Tris-HCl, 45 mM NaCl, 1 mM MgCl_2_, 1.6% Glycerol, 133 nM GDP, 2 *µ*M [γ-^32^P] GTP, 50 *µ*M AppNHp, 67 *µ*M β-ME, 67 *µ*M AEBSF, in the presence or absence of 8 nM FLAG-Sec12 (1–386 aa)-His_6_, 15 nM HA-Sec23A/FLAG-Sec24D, and 200 nM Sar1, and indicated concentrations of His_6_-cTAGE5 (61–804 aa)-FLAG and Sec13/31. His_6_-GST or His_6_-cTAGE5 (651–804 aa)-GST wild-type or R757G or 4PA, FLAG-Sec12 (1–386 aa)-His_6_ and liposome were preincubated in a buffer consisting of 20 mM Hepes-KOH, pH 7.4, 160 mM KOAc, 1 mM MgCl_2_ for 2h at 4°C Then, Sar1 GTP hydrolysis were initiated in a buffer consisting of 25 *µ*g/ml of liposome, 20 mM HEPES-KOH (pH 7.4), 100 mM KOAc, 8 mM Tris-HCl, 60 mM NaCl, 1 mM MgCl_2_, 1.6% Glycerol, 133 nM GDP, 2 *µ*M [γ-^32^P] GTP, 50 *µ*M AppNHp, 67 *µ*M β-ME, 67 *µ*M AEBSF, 8 nM FLAG-Sec12 (1–386 aa)-His_6_, 15 nM HA-Sec23A/FLAG-Sec24D, and 200 nM Sar1, and indicated concentrations of His_6_-GST, His_6_-cTAGE5 (651–804 aa)-GST wild-type, R757G, 4PA. The reaction mixtures were incubated for 1 h at 30°C, and quenched by adding 750 *µ*L of ice-cold 5%(w/v) Norit SX-Plus (Wako, Japan) in 50 mM NaH_2_PO_4_, then centrifuged at 9,000 g for 15 min at 4°C 300 *µ*L of supernatants were mixed with 1 mL of Clear-sol I (Nacalai-tesque, Japan) and free ^32^Pi was measured.

### Random mutagenesis screening

Random mutagenesis was essentially performed as described previously (Cadwell and Joyce, 1992). Error-prone PCR was performed with Titanium Taq polymerase (Takara, Japan) in the presence of 0.64 mM MnCl_2_ with unbalanced ratio of nucleotides (0.2 mM dATP, 1 mM dTTP, 1 mM dGTP, 1 mM dCTP). PCR products were cloned into pGADT7 for following yeast two-hybris analysis. AH109 yeast strain was transformed with pGBKT7 and pGADT7 vectors and plated on tryptophan- and leucine-deficient plate. The colonies were re-plated onto tryptophan-, leucine-, histidine-, and adenine-deficient plate. The colonies, which failed to grow on the tryptophan-, leucine-, histidine-, and adenine-deficient plate were picked and lysed by zymolyase for sequence analysis.

### *In vitro* binding assay

GST-cTAGE5 (651–804 aa)-His_6_ wild-type, GST-cTAGE5 (651–804 aa)-His_6_ R757G, GST-cTAGE5 (651–804 aa)-His_6_ 4PA expressed in *Escherichia* were purified with glutathione sepharose followed by elution with glutathione. Eluates were then purified with Ni sepharose 6 Fast Flow and eluted with imidazole. The buffers were exchanged with TBS/0.05% Lubrol-PX by desalting columns. *In vitro* binding assay was essentially performed as described previously (Maeda *et al.*, 2017). In brief, GST, GST-tagged cTAGE5-PRD (651–804 aa)-His_6_ constructs were conjugated to glutathione sepharose and incubated with FLAG-Sec23A. Beads were washed with TBS/0.05% Lubrol-PX for five times followed by elution with glutathione.

### Immunoprecipitation and Western blotting

The experiments were performed essentially as described previously (Maeda *et al.*, 2017). In brief, Cells extracted with extraction buffer (20 mM Tris-HCl (pH 7.4), 100 mM NaCl, 1 mM EDTA, 1% Triton X-100, and protease inhibitors) were centrifuged at 100,000 × *g* for 30 min at 4°C. Cell lysates were immunoprecipitated with FLAG M2 Agarose beads (Sigma-Aldrich). The beads were washed five times with TBS/0.1% Triton X-100 and processed for sample preparation.

### Immunofluorescence microscopy

Immunofluorescence microscopy analysis was performed as described previously (Maeda *et al.*, 2017). Cells grown on coverslips were washed with phosphate-buffered saline (PBS), fixed with methanol (6 min at −20°C), and then washed with PBS and blocked in blocking solution (5% bovine serum albumin in PBS with 0.1% Triton X-100 for 15 min). After blocking, cells were stained with primary antibody for 1 h, followed by incubation with Alexa Fluor–conjugated secondary antibodies for 1 h at room temperature. Images were acquired with confocal laser scanning microscopy (LSM700; Plan-Apochromat 63×/1.40 numerical aperture [NA] oil immersion objective lens; Carl Zeiss, Oberkochen, Germany). The acquired images were processed with Zen 2009 software (Carl Zeiss). All imaging was performed at room temperature.

### Quantification of collagen VII staining

Quantification of collagen VII accumulation was essentially performed as described previously (Tanabe *et al.*, 2016). Stained cells were analyzed by Zeiss Axio Imager M1 microscopy (EC Plan-Neofluar 40×/ 0.75 NA objective lens) and processed with AxioVision software (Carl Zeiss). Area calculation and intensity scanning were done by ImageJ software (National Institutes of Health, Bethesda, MD).

## Acknowledgements

We thank Dr. Lynn Sakai for providing us NP185 antibody. We also thank members of the Katada lab for valuable discussions. This work was supported in part by research grants from Japan Society for the Promotion of Science (JSPS) (grant numbers 17J07885 to M.M.; 23229001 to T.K., 26440046, 17H03651 to K.S.).

---

**Supplemental Figure 1.**
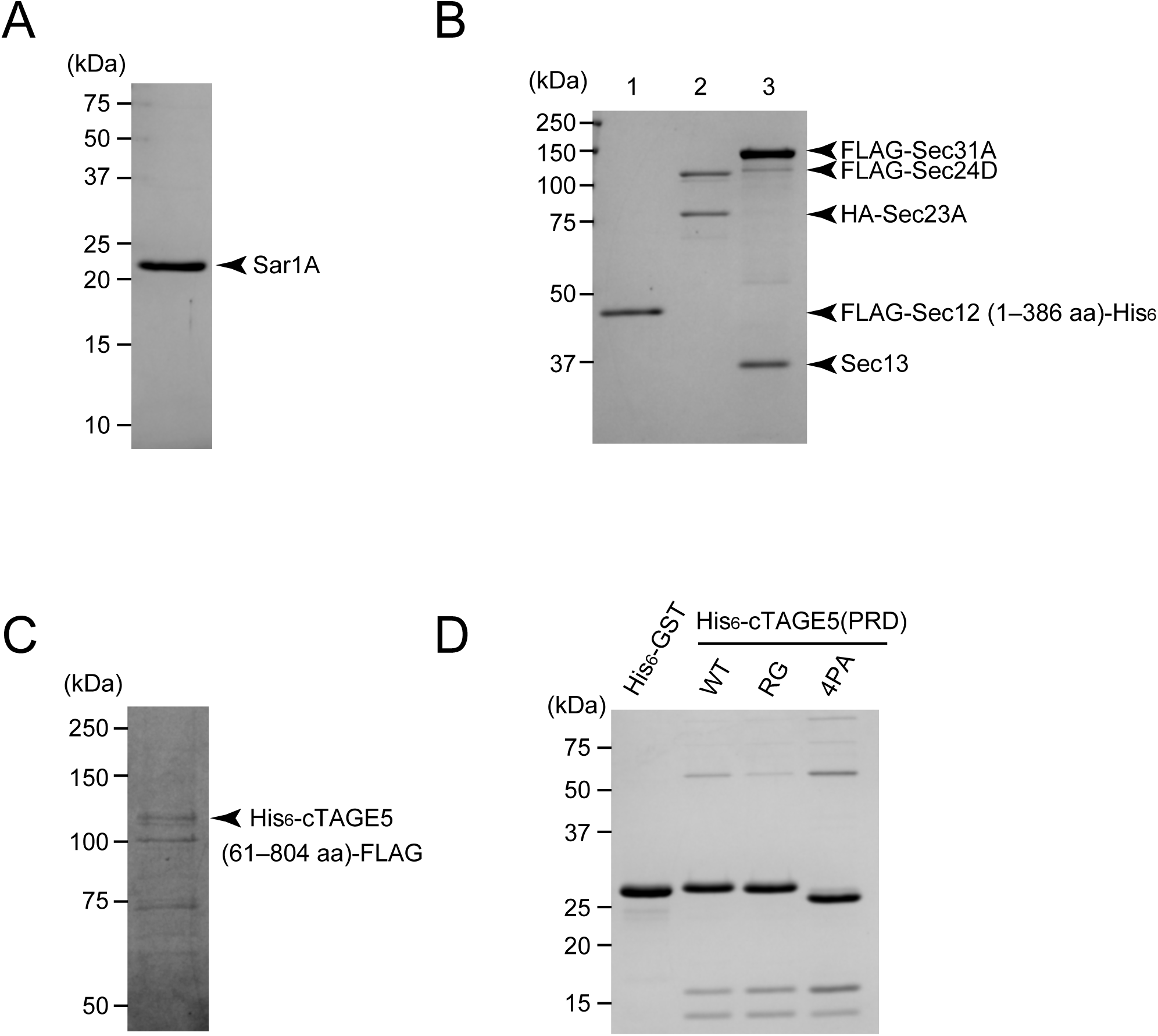
Recombinant human proteins used in the GTPase hydrolysis assay were resolved by SDS-PAGE followed by CBB staining. (A) 1.0 *µ*g Sar1A. (B) 0.6 *µ*g FLAG-Sec12 (1–386 aa)-His_6_ (lane 1), 0.43 *µ*g HA-Sec23A/0.6 *µ*g FLAG-Sec24D (lane 2), 0.54 *µ*g Sec13/2.0 *µ*g FLAG-Sec31A (lane 3). (C) 10 ng His_6_-cTAGE5 (61–804 aa)-FLAG. (D) 2.6 *µ*g His_6_-GST, 2.25 *µ*g His_6_-cTAGE5 (PRD) WT, RG, 4PA.

**Supplemental Figure 2.**
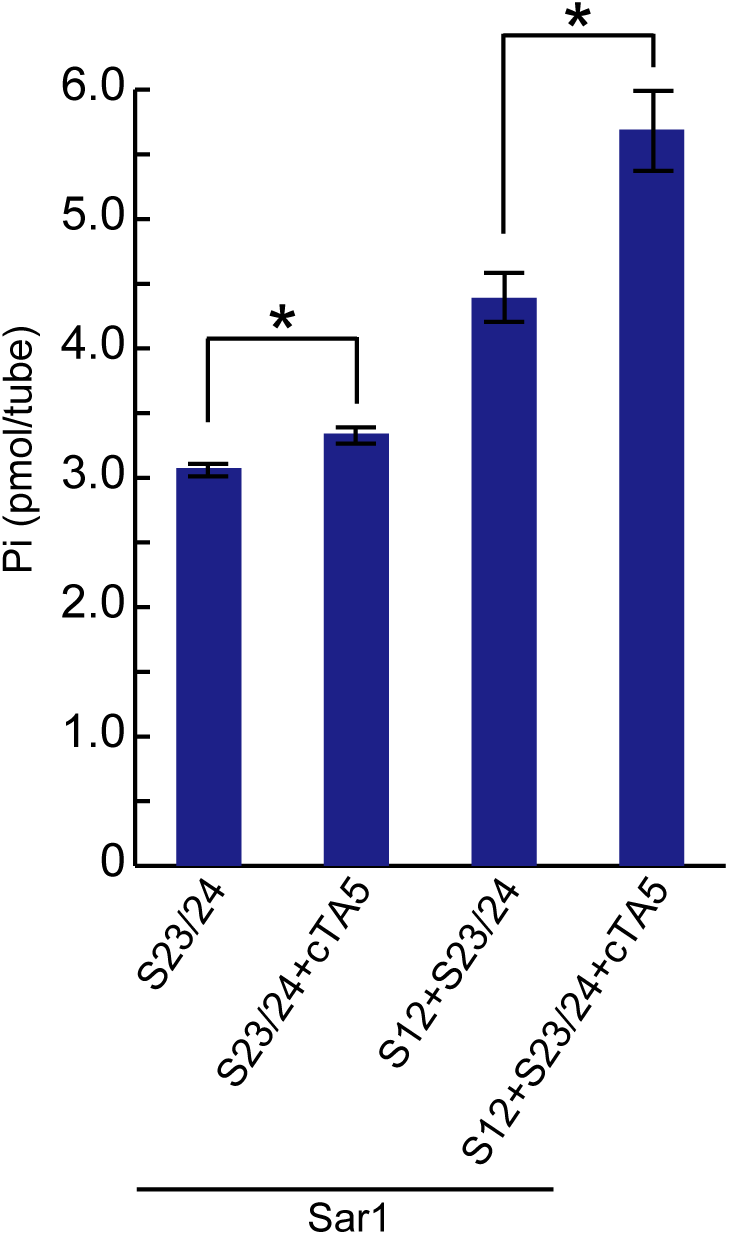
200 nM Sar1 and 15 nM HA-Sec23A/FLAG-Sec24D were incubated with a buffer consisting 25 *µ*g/ml Ni liposome (20%), 20 mM HEPES-KOH (pH 7.4), 100 mM KOAc, 6 mM Tris-HCl, 45 mM NaCl, 1 mM MgCl_2_, 1.6% Glycerol, 133 nM GDP, 2 *µ*M [γ-^32^P] GTP, 50 *µ*M AppNHp, 67 *µ*M β-ME, 67 *µ*M AEBSF, in the presence or absence of 8 nM FLAG-Sec12 (1–386 aa)-His_6_, and 3.6 nM cTAGE5 for 1 h at 30°C. The amount of free ^32^Pi was quantified, *n* = 6. *p<0.05. Error bars represent means ± SEM.

**Supplemental Figure 3.**
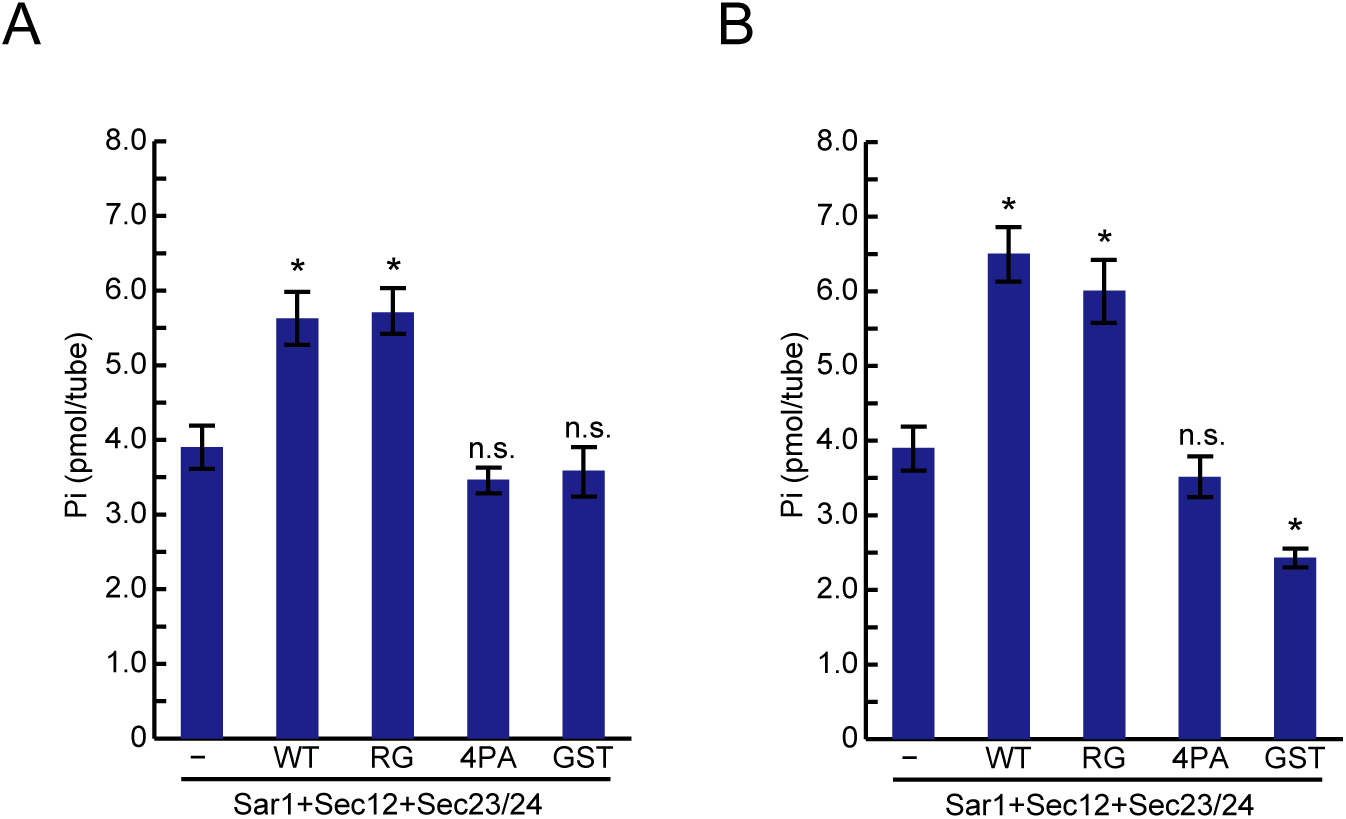
200 nM Sar1 was incubated with a buffer consisting 25 *µ*g/ml Ni liposome (10%), 20 mM HEPES-KOH (pH 7.4), 100 mM KOAc, 8 mM Tris-HCl, 60 mM NaCl, 1 mM MgCl_2_, 1.6% Glycerol, 133 nM GDP, 2 *µ*M [γ-^32^P] GTP, 50 *µ*M AppNHp, 67 *µ*M β-ME, 67 *µ*M AEBSF, 8 nM FLAG-Sec12 (1–386 aa)-His_6_, 15 nM HA-Sec23A/FLAG-Sec24D and 400 nM (A) or 800 nM (B) of His_6_-GST, His_6_-cTAGE5 (651–804 aa)-GST wild-type, R757G, 4PA for 1 h at 30°C. The amount of free ^32^Pi was quantified, *n* = 5. *p<0.05. n.s., not significant compared with His_6_-GST. Error bars represent means ± SEM.

